# Molecular recognition of an acyl-peptide hormone and activation of ghrelin receptor

**DOI:** 10.1101/2021.06.09.447478

**Authors:** Yue Wang, Shimeng Guo, Youwen Zhuang, Ying Yun, Peiyu Xu, Xinheng He, Jia Guo, Wanchao Yin, H. Eric Xu, Xin Xie, Yi Jiang

**Affiliations:** CAS Key Laboratory of Receptor Research, Center for Structure and Function of Drug Targets, Shanghai Institute of Materia Medica, Chinese Academy of Sciences, Shanghai 201203, China; University of Chinese Academy of Sciences, Beijing 100049, China; School of Chinese Materia Medica, Nanjing University of Chinese Medicine, Nanjing 210046, China; CAS Key Laboratory of Receptor Research, National Center for Drug Screening, Shanghai Institute of Materia Medica, Chinese Academy of Sciences, Shanghai 201203, China; School of Life Science and Technology, ShanghaiTech University, Shanghai 201210, China; School of Pharmaceutical Science and Technology, Hangzhou Institute for Advanced Study, UCAS, Hangzhou 310024

## Abstract

Ghrelin, also called “the hunger hormone”, is a gastric peptide hormone that regulates food intake, body weight, as well as taste sensation, reward cognition, learning and memory. One unique feature of ghrelin is its acylation, primarily with an octanoic acid, which is essential for its binding and activation of the ghrelin receptor, a G protein-coupled receptor. The multifaceted roles of ghrelin make ghrelin receptor a highly attractive drug target for growth retardation, obesity, and metabolic disorders. Here we present two cryo-electron microscopy structures of G_q_-coupled ghrelin receptor bound to ghrelin and a synthetic agonist, GHRP-6. Analysis of these two structures reveals a unique binding pocket for the octanoyl group, which guides the correct positioning of the peptide to initiate the receptor activation. Together with mutational and functional data, our structures define the rules for recognition of the acylated peptide hormone and activation of ghrelin receptor, and provide structural templates to facilitate drug design targeting ghrelin receptor.

## Introduction

Food intake is one of the most fundamental processes required for sustaining human life. It is primarily regulated by two endogenous hormones with opposite physiological functions: leptin, the energy surfeit hormone, and ghrelin, the hunger hormone, both of which are involved in controlling energy balance and obesity. Ghrelin is an orexigenic peptide hormone secreted from stomach in response to fasting situations and stimulates the ghrelin receptor in the brain to initiate appetite ^1–4^. One unique feature of ghrelin is the fatty acid modification, with its third amino acid Ser^3^ being modified with an octanoyl group ^1^, catalyzed by ghrelin O-acyltransferase (GOAT) ^5,6^. Although less than 10% of ghrelin is acylated in the blood ^7^, this acyl-modification is essential for its activity. Both ghrelin and synthesized growth hormone secretagogues show potent growth hormone-releasing activity and serve as potential candidates for the treatment of growth hormone deficiency (GHD) ^8^. The growth hormone-releasing activity also makes these hormones attractive performance-enhancing substances, whose usages are banned by the World Anti-Doping Agency in competitive sports ^9^.

The pleiotropic functions of ghrelin are mediated through ghrelin receptor, also known as the growth hormone secretagogue receptor, which was first identified in the pituitary gland and the hypothalamus. As a G protein-coupled receptor (GPCR), ghrelin receptor couples to G_q_ protein and modulates diverse physiological processes upon binding to ghrelin and other synthetic agonists ^10^. LEAP2, an intestinally derived hormone, is identified as an endogenous antagonist of ghrelin receptor, which fine-tunes ghrelin action via an endogenous counter-regulatory mechanism ^11^. Ghrelin receptor is characterized by its high basal activity, with approximately 50% of its maximal capacity in the absence of a ligand ^12,13^. This high level of basal activity may serve as a “signaling set point” to counterbalance the inhibitory input from leptin and insulin in appetite regulation ^12^. Several naturally occurring mutations of *ghrelin receptor*, such as A204E and F279L, decrease the basal activity of the receptor and have been found to associate with obesity, diabetes, and short stature ^14^, which led to the idea of ghrelin receptor being an attractive therapeutic target for these diseases. However, there are only two orally-active synthetic agonists, pralmorelin and macimorelin, been approved as diagnostic agents for GHD to date.

Extensive efforts have been devoted to examining the structural basis for the potential binding sites of ghrelin and synthetic agonists ^13,15–19^ and the basal activity of ghrelin receptor ^20–22^. Nevertheless, compared to leptin and its receptor, which structures are known ^23,24^, much less are known about the structures of ghrelin and ghrelin-bound receptor. In this study, we reported two cryo-electron microscopy (cryo-EM) structures of the active ghrelin receptor–G_q_ complexes bound to ghrelin and GHRP-6, respectively.

## Results and Discussion

### Structural determination

We fused thermostabilized BRIL at the N-terminus of ghrelin receptor and applied the NanoBiT tethering strategy to improve complex stability and homogeneity (Extended Data Fig. 1a, b) ^25^. An engineered Gα_q_ was based on the mini-Gα_s_ scaffold with its N-terminus replaced by corresponding sequences of Gα_i1_ to facilitate the binding of scFv16 (Extended Data Fig. 1c), an analogous approach had been used to obtain structures of the G_q_-bound 5-HT_2A_ receptor ^26^ and G_11_-bound M1 receptor ^27^. Unless otherwise specified, G_q_ refers to the engineered G_q_, which is used for further structure study. Ghrelin receptor was co-expressed with Gα_q_ and Gβγ, and incubated with ghrelin in the presence of Nb35 to stabilize the receptor-G protein complex ^28^, allowing the efficient assembly of the ghrelin receptor–G_q_ complex. The scFv16 was additionally added to assemble the GHRP-6–ghrelin receptor–G_q_–scFv16 complex (Extended Data Fig. 2d).

The complex structures of the G_q_-coupled ghrelin receptor bound to ghrelin and GHRP-6 were determined by cryo-EM to the resolutions of 2.9 Å and 3.2 Å, respectively (Fig. 1a-d, Extended Data Fig. 2, Extended Data Table 1). For both ghrelin receptor–G_q_ complexes, the majority of the amino acid side-chains of receptor and G_q_ protein were well-resolved in the final models, which are refined against the EM density map with excellent geometry. Both peptides, ghrelin (Gly^1P^-Arg^15P^) with octanoylated modification and GHRP-6, were clearly identified, thus providing reliable models for the mechanistic explanation of peptide recognition and activation of ghrelin receptor (Extended Data Fig. 3).

**Fig. 1.**
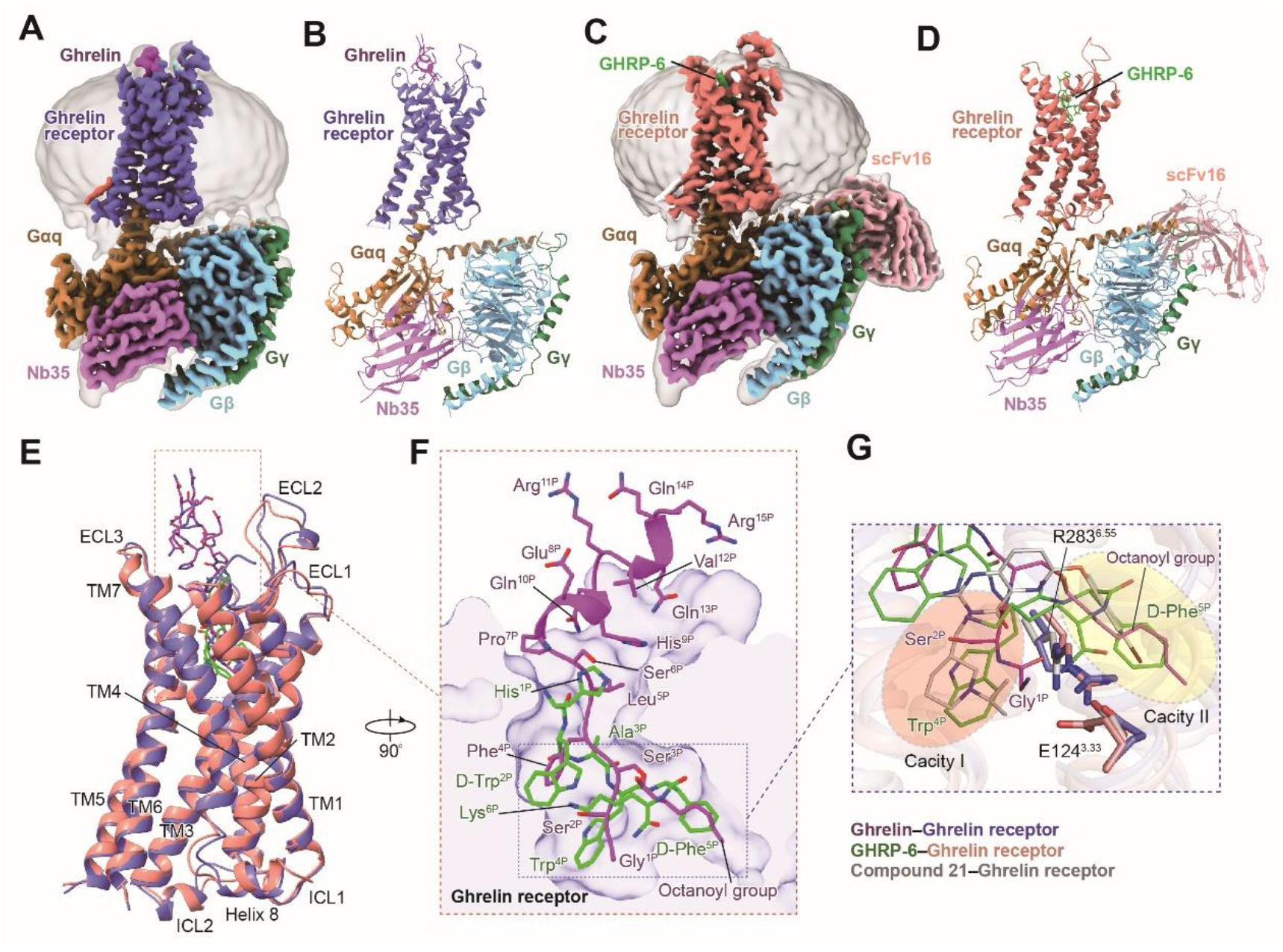
Cryo-EM structures of the G_q_-coupled ghrelin receptor bound to ghrelin and GHRP-6. **a, b**, Orthogonal views of the density map (**a**) and model (**b**) for the ghrelin–ghrelin receptor–G_q_–Nb35 complex. The density map is shown at 0.104 threshold. **c, d**, Orthogonal views of the density map (**c**) and model (**d**) for the GHRP-6–ghrelin receptor–G_q_–Nb35–scFv16 complex. The density map is shown at 0.1 threshold. **e**, Structural superposition of ghrelin-bound and GHRP-6-bound ghrelin receptors. **f**, Binding poses of ghrelin and GHRP-6. Two peptides occupy a similar orthosteric binding pocket with opposite orientation. **g**, Binding pocket of ghrelin receptor is bifurcated into two cavities by a salt bridge between E124^3.33^ and R283^6.5518^. Salmon oval, cavity I; yellow oval, cavity II. Ghrelin is shown in magenta, ghrelin-bound ghrelin receptor in slate blue. GHRP-6 is displayed in green, and GHRP-6 bound ghrelin receptor in salmon. Compound 21-bound ghrelin receptor (PDB: 6KO5) is colored grey. The G_q_ heterotrimer is colored by subunits. Gα_q_: peru, Gβ: light sky blue, Gγ: sea green, Nb35: orchid, scFv16: light pink.

### Overall structures of G_q_-coupled ghrelin receptor bound to ghrelin and GHRP-6

Both ghrelin–ghrelin receptor–G_q_ and GHRP-6–ghrelin receptor–G_q_ complexes present canonical folds of seven transmembrane segments with the TMD of the receptors surrounded by an annular detergent micelle mimicking the natural phospholipid bilayer (Fig. 1a-d). Within the micelle, two cholesterols are clearly visible and hydrophobically bind around the helix bundles of both ghrelin receptor complexes. Both complexes display highly identical overall conformations with the root mean square deviation (RMSD) of 0.8 Å for entire complexes and 0.6 Å for ghrelin receptors (Fig. 1e). Ghrelin and GHRP-6 occupy the same orthosteric ligand-binding pocket of ghrelin receptor, comprising of all TM helices and extracellular loops (ECLs) except TM1 and ECL1 (Fig. 1e, Extended Data Fig. 4). The N-terminus of ghrelin inserts deep in the helix bundle. Conversely, GHRP-6 adopts an upside-down binding mode relative to ghrelin, with its C-terminus inserting into the helix bundle and its N-terminus facing the extracellular vestibule (Fig. 1f). Additionally, GHRP-6 is largely overlaid with the first six amino acids fragment of ghrelin (Fig. 1f). Recently, the structure of ghrelin receptor bound to an antagonist compound 21 revealed a characteristic feature that the binding pocket is bifurcated into two cavities by a salt bridge between E124^3.33^ and R283^6.5518^. Both ghrelin and GHRP-6 adopt similar binding modes and are buried in two identical cavities relative to compound 21, revealing a conserved binding pose for both peptidic agonists and antagonists (Fig. 1g).

### Molecular basis for recognition of ghrelin by ghrelin receptor

The N-terminal amino acids from Gly^1P^ to Pro^7P^ of ghrelin occupied nearly the entire receptor TMD binding pocket (Fig. 2a). The peptide fragment from Glu^8P^ to Arg^15P^ is enriched with polar and charged amino acids and adopts an α-helical conformation, which sits above the orthosteric pocket and interacts with the solvent (Fig. 2a).

**Fig. 2.**
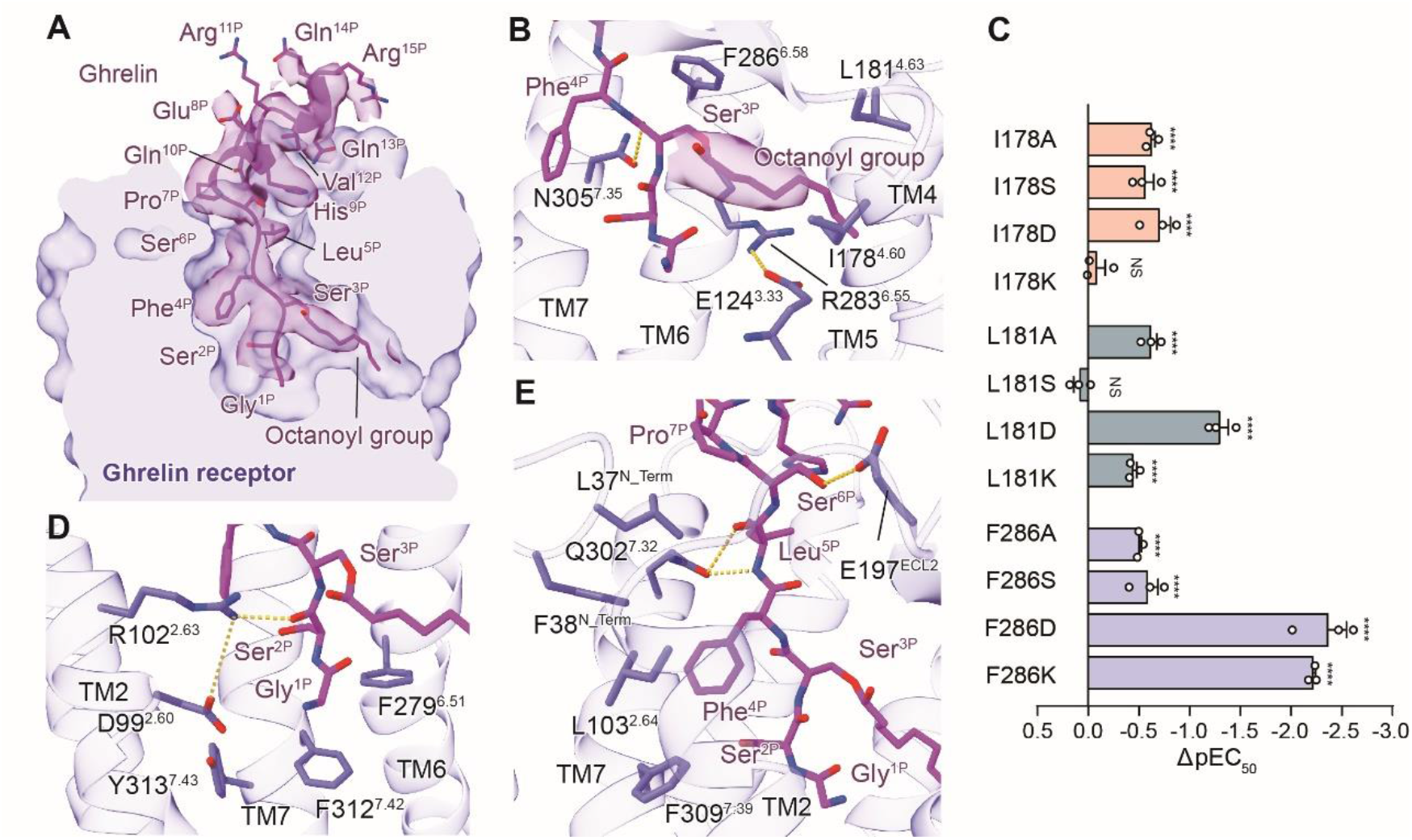
The ghrelin-binding pocket of ghrelin receptor. **a**, Cross-section of the ghrelin-binding pocket in ghrelin receptor. The cryo-EM density of ghrelin is highlighted. Ghrelin is shown in cartoon presentation. Side chains of the residues are displayed as sticks. **b**, The binding pocket for octanoyl group. The cryo-EM density of the octanoyl group is shown. The hydrogen bonds are depicted as yellow dashed lines. **c**, Effects of mutations of residues in the octanoyl group-binding pocket on calcium response. ΔpEC50 represents the difference between pEC_50_ values of the mutant ghrelin receptor and the wild-type (WT) receptor. Data are presented as mean ± S.E.M. of three independent experiments performed in technical triplicate. All data were analyzed by two-side, one-way ANOVA with Tukey’s test. \*\*\*\**P*<0.0001 *vs*. WT receptor, NS, no significant difference. **d, e**, Detailed interactions of ghrelin (Gly^1P^-Pro^7P^) with residues in ghrelin binding pocket. Ghrelin is shown in magenta, and ghrelin receptor in slate blue.

The clearly visible map allows us to locate Ser^3P^ and its octanoyl group accurately. A density connects the side-chain of Ser^3P^ and stretches horizontally toward the gap between TM4 and TM5, occupying cavity II of the binding pocket. Five carbons of the eight-carbon fatty acid modification can be placed in the density (Fig. 2b). F286^6.58^ covers the side-chain of Ser^3P^, while the fatty acid chain of ghrelin forms hydrophobic contacts with I178^4.60^ and L181^4.63^ (Figs. 2b, 3e). Most substitutions of I178^4.60^, L181^4.63^, and F286^6.58^ with alanine or polar and charged amino acids (Ser, Asp, or Lys) display dramatically diminished receptor activation compared with wild-type (WT) ghrelin receptor (Fig. 2c, Extended Data Table 2). It is worth noting that F286^6.58^ makes a greater contribution to ghrelin’s activity. Mutating F286^6.58^ to Asp produces an over 250-fold decreased activity of ghrelin (Fig. 2c), while mutating L178^4.60^ or I181^4.63^ to Asp diminishes the activities by only ~5-fold and 20-fold, respectively (Fig. 2c, Extended Data Table 2). These findings highlight the critical role of the potent receptor hydrophobic environment for binding of the octanoyl group. These results are consistent with the previous report that replacement of Ser^3P^ by a charged amino acid (Lys) or small hydrophobic residues (Val, Leu, or Ile) significantly decreased ghrelin’s activity ^29^. Conversely, replacing Ser^3P^ with aromatic amino acids, such as Trp or β-Nal (2-naphtylalanine), preserved ghrelin’s activity ^29^.

**Fig. 3.**
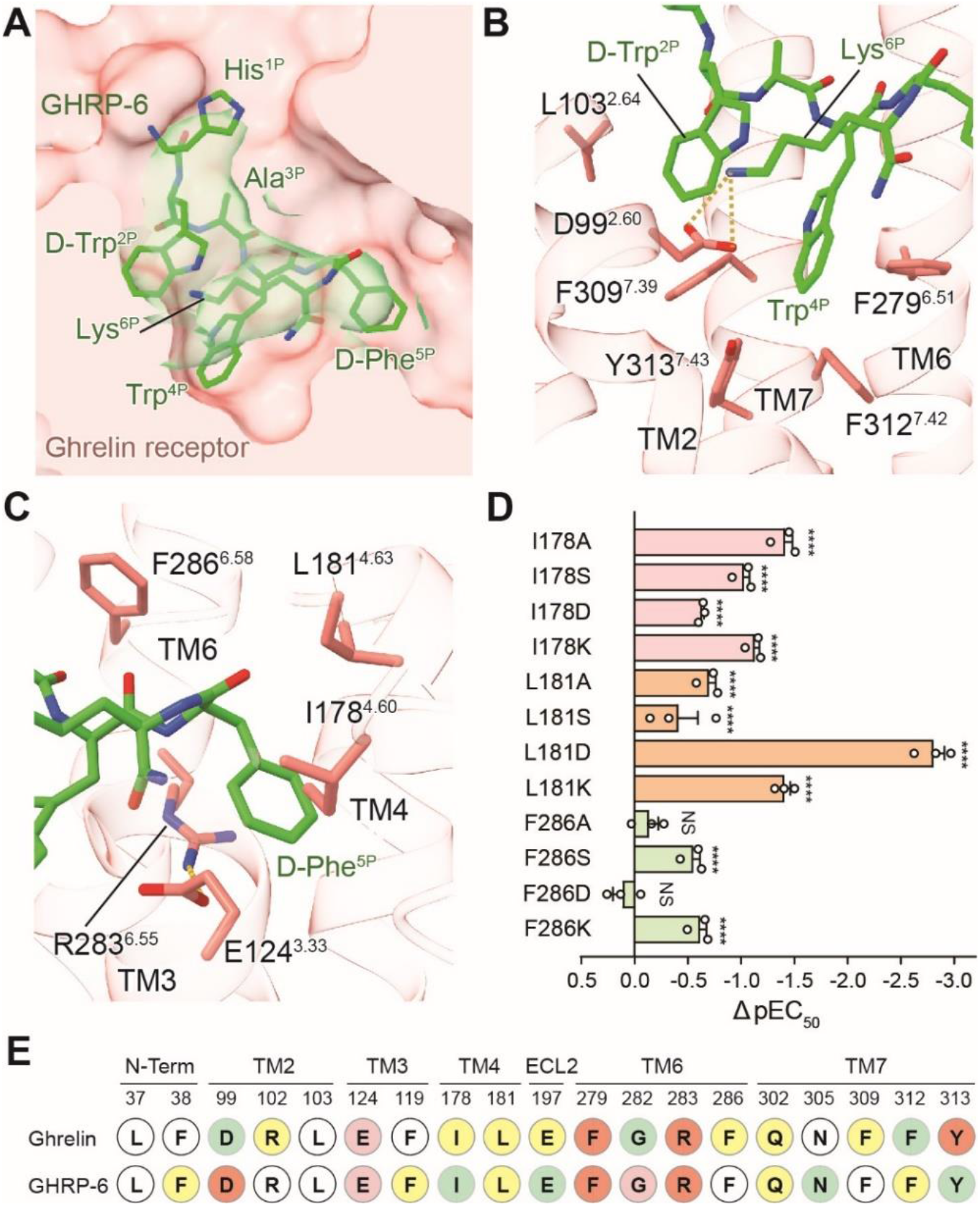
The GHRP-6-binding pocket of ghrelin receptor. **a**, Cross-section of the GHRP-6-binding pocket in ghrelin receptor. The cryo-EM density of GHRP-6 is highlighted. GHRP-6 is shown in a cartoon presentation. Side-chains of the residues are displayed as sticks. **b**, Detailed interactions of GHRP-6 (His^1P^-Lys^6P^) with residues in GHRP-6 binding pocket. The hydrogen bond is depicted as a yellow dashed line. **c**, The binding site of D-Phe^5P^. D-Phe^5P^ is highly overlaid with the octanoyl group of ghrelin. **d**, Effects of mutations of residues in D-Phe^5P^ binding pocket on calcium response. ΔpEC50 represents the difference between pEC_50_ values of the mutant ghrelin receptor and the wild-type (WT) receptor. Data are presented as mean ± S.E.M. of three independent experiments performed in technical triplicate. All data were analyzed by two-side, one-way ANOVA with Tukey’s test. \*\*\*\**P*<0.0001 *vs*. WT receptor, NS, no significant difference. GHRP-6 is shown in green, and ghrelin receptor in salmon. **e**, Interaction residues in the ligand binding pocket of the ghrelin–ghrelin receptor–G_q_ and the GHRP-6–ghrelin receptor–G_q_ complexes. Compared with WT receptor, the alanine replacement of residues showed comparable impacts on peptide’s activity are indicated by white circles. The alanine mutation of residues showed 2-10-fold, 10-100-fold, 100-1000-fold, and over 1000-fold decreased peptide’s activity are indicated by yellow, green, salmon, and red circles, respectively.

Besides these hydrophobic residues surrounding the octanoyl group, other pocket residues also make substantial contributions to ghrelin binding and receptor activation. R283^6.55^ forms a stabilizing salt bridge with E124^3.33^ to lock TM3 and TM6 (Fig. 2b). Alanine mutations of E124^3.33^ and R283^6.55^ nearly abolished the binding of ghrelin-A2 and the ghrelin-induced receptor activity, indicating the critical role of the salt bridge in maintaining the integrity of the bifurcated binding pockets, and thus affecting ghrelin binding and receptor activation (Extended Data Figs. 5, 6b, Extended Data Table 2). Gly^1P^ and Ser^2P^ locate in cavity I of the ghrelin receptor binding pocket (Fig. 2d). The main chain CO group of Ser^2P^ H-bonds with the side-chain of R102^2.63^, which is critical for ghrelin binding (Fig. 2d, Extended Data Figs. 5, 6, Extended Data Table 2). Meanwhile, the N-terminal amino acid Gly^1P^ engages hydrophobic interactions with F279^6.51^ and F312^7.42^, which sit at the bottom of the binding pocket (Fig. 2d). It should be noted that alanine mutations of F279^6.51^ and F312^7.42^ partly maintain the binding of ghrelin but remarkably diminish its activity, indicating that these phenylalanines are mainly responsible for signal transmission (Fig. 2d, Extended Data Figs. 5, 6b, Extended Data Table 2). This finding suggests that the insertion of the octanoyl group into cavity II not only contributes to the binding of the ligand, but also helps to orient the N-terminus of ghrelin to cavity I, which is critical in receptor activation.

Additionally, Phe^4P^ is surrounded by hydrophobic residues, including L37^N_Term^, F38^N_Term^, L103^2.64^, and F309^7.39^, all contributing to ghrelin binding and its activity (Fig. 2e; Extended Data Figs. 5, 6b, Extended Data Table 2). Furthermore, Leu^5P^ forms an H-bond with Q302^7.32^ via its mainchain, while Ser^6P^ connects to ECL2 by forming an H-bond with E197^ECL2^, both residues contribute to ghrelin binding (Figs. 2e, 3e, Extended Data Figs. 5, 6b; Extended Data Table 2). These detailed structural analyses provide insights to understand the recognition mechanism of the acyl-modified ghrelin by ghrelin receptor.

### Molecular basis for recognition of GHRP-6 by ghrelin receptor

GHRP-6, a synthetic peptidic growth hormone secretagogue derived from met-enkephalin, shows no sequence homology with ghrelin ^30^. It is buried in the same orthosteric site of ghrelin receptor and displays comparable potency for receptor activation (Fig. 3a, Extended Data Fig. 6a). GHRP-6 adopts a similar binding pose with Gly^1P^-Ser^6P^ of ghrelin (Fig. 1f). Cavity I accommodates Trp^4P^ of GHRP-6 and offers a more extensive hydrophobic environment comprising F279^6.51^, F309^7.39^, F312^7.42^, and Y313^7.43^, of which F279^6.51^ and Y313^7.43^ are closely related to the activity of GHRP-6 (Figs. 1g, 3b, 3e, Extended Data Figs.6c, Extended Data Table 2). D-Phe^5P^ occupies cavity II and is highly overlaid with the entire octanoyl group of ghrelin (Figs. 1f, 3c). Besides its hydrophobic contacts with I178^4.60^, L181^4.63^, and F286^6.58^, D-Phe^5P^ forms an extra cation-π interaction with R283^6.55^ relative to ghrelin (Fig. 3c). Although alanine substitutions of I178^4.60^, L181^4.63^, and F286^6.58^ all significantly impair activities of ghrelin and GHRP-6, these residues make distinct extents contributions. In contrast to ghrelin, replacement of I178^4.60^ or L181^4.63^ by alanine demonstrates a more remarkable decreased GHRP-6’s activity than F286^6.58^ (Fig. 3d, Extended Data Table 2).

In addition, D-Trp^2P^ forms an edge-to-face packing with Trp^4P^ and establishes a stabilizing intramolecular hydrophobic network with the side-chain of Lys^6P^ (Fig. 3b). A previous study reported that growth hormone secretagogue metabolites without Lys^6P^ abolished its ghrelin receptor binding capacity ^8^. This finding is consistent with our observation that the side-chain of Lys^6P^ points to TM2 and forms a stabilizing salt bridge with D99^2.60^, which is closely related to ghrelin receptor activation (Figs. 3b, 3e, Extended Data 6c, Extended Data Table 2). Our structural finding highlights the significance of Lys^6P^ on GHRP-6’s activity. Together, these results reveal the recognition mechanism of GHRP-6 and diversify the peptide-binding mode for ghrelin receptor.

### Activation mechanism of ghrelin receptor

Structural comparison of two G_q_-coupled ghrelin receptors with other G_q/11_-coupled class A GPCRs reveals similar receptor conformations. TM6 and TM7 of ghrelin receptor adopt nearly identical conformations to G_q_-coupled 5-HT_2A_ receptor and G_11_-coupled M1 receptor (Extended Data Fig. 7a, b). Unlike M1R, the cytoplasmic end of TM5 of ghrelin receptor does not show an inward movement (Extended Data Fig. 7a). In addition, the Gα α5 helix displays slightly shifts and the distal end of Gα_q_ α5 helix shows more notable shifts across these receptor complexes (Extended Data Fig. 7c). Furthermore, structural comparison of two G_q_-coupled ghrelin receptors with the antagonist-bound ghrelin receptor (PDB: 6KO5) ^18^ supports the contention that these two complexes are indeed in the active state (Fig. 4a, b). These two ghrelin receptor complexes display pronounced outward displacements of TM6 cytoplasmic end (~8 Å, measured at Cα of H258^6.30^), the hallmark of GPCR activation, and ~3 Å inward shift of cytoplasmic end of TM7 (measured at Cα of Y323^7.53^) (Fig. 4a, b).

**Fig. 4.**
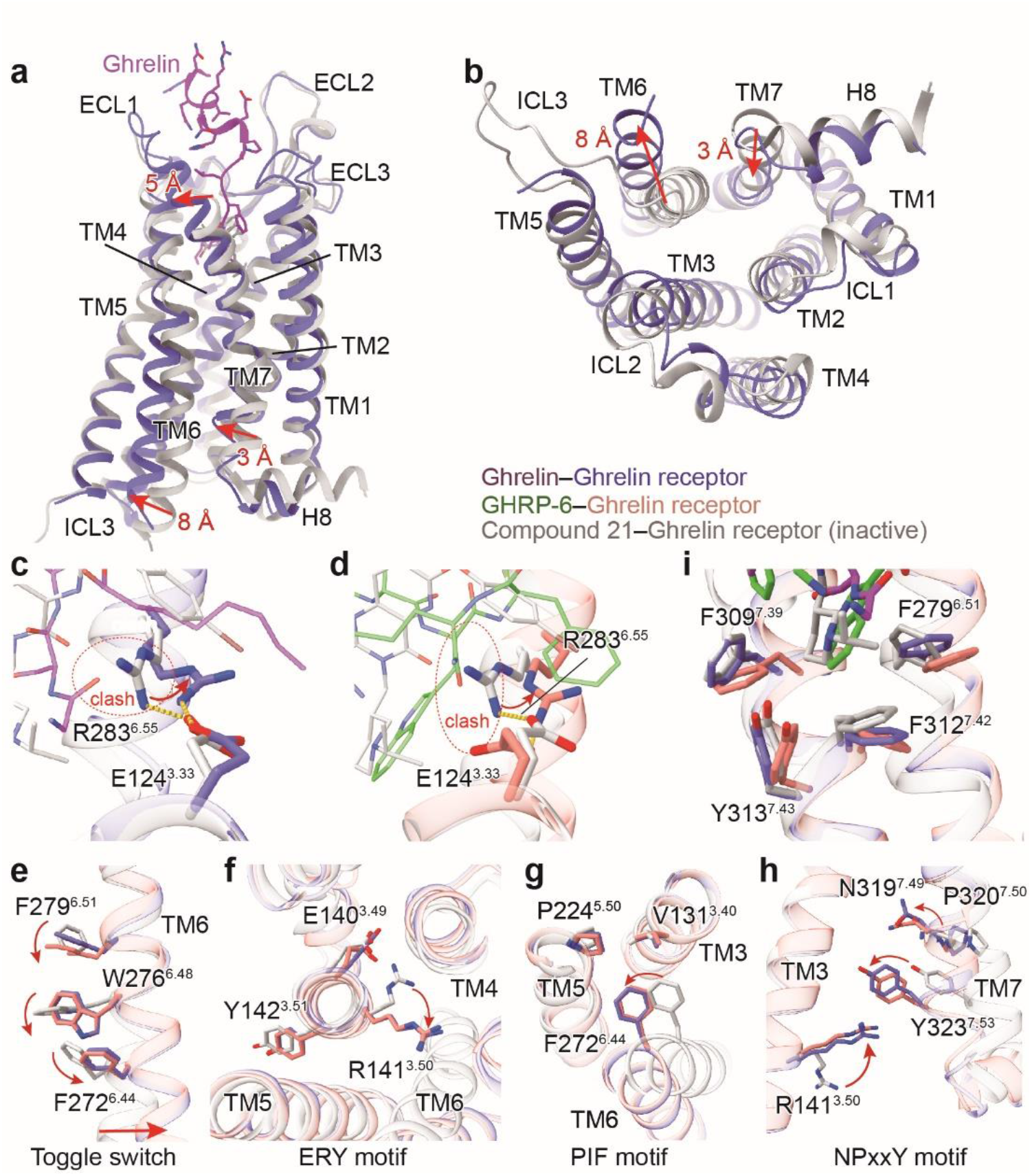
Activation mechanism of ghrelin receptor. **a, b**, Structural superposition of the active and antagonist-bound ghrelin receptor. (**a**) side view; (**b**) cytoplasmic view. **c, d**, Conformational changes of the salt bridge comprising of E124^3.33^ and R283^6.55^ upon peptide activation. Ghrelin and GHRP-6 push the side-chain of R283^6.55^ to swing away from the receptor helical core due to steric clash. The swing orientations of E124^3.33^ and R283^6.55^ were indicated by red arrows. The clashes were highlighted as red oval dashed lines. **e-h**, Conformational changes of the conserved “micro-switches” upon receptor activation. (**e**) Toggle switch, (**f**) ERY motif, (**g**) PIF motif, (**h**) NPxxY motif. The outward displacement of TM6 of the active receptor is shown as a red arrow (**e**). The conformational changes of residue side chains are shown as red arrows upon receptor activation. The hydrophobic cluster comprising F279^6.51^, F309^7.39^, F312^7.42^, and Y313^7.43^ stabilizes the inter-helical hydrophobic contacts between TM6 and TM7. Ghrelin is shown in magenta, ghrelin-bound ghrelin receptor in slate blue. GHRP-6 is displayed in green, and GHRP-6 bound ghrelin receptor in salmon. The compound 21-bound ghrelin receptor (PDB: 6KO5), is colored in grey.

Although ghrelin and GHRP-6 differ in their chemical scaffolds, they induce the activation of ghrelin receptor through similar conformational changes in TM6 as well as TM7. Compared to the antagonist-bound receptor, the extracellular end of TM7 of these two active receptors laterally move ~5 Å towards TM6 (measured at Cα of Q302^7.32^), presenting a unique TM7 conformational change across class A GPCRs solved to date (Fig. 4a, Extended Data Fig. 8). The notable movement of the extracellular end of TM7 may translate to the inward shift of its cytoplasmic end. Furthermore, structural comparison of both active and antagonist-bound receptors supports that R283^6.55^ is the determinant for peptide-induced ghrelin receptor activation. The salt bridge between R283^6.55^ and E124^3.33^ exists in both antagonist-bound and active ghrelin receptor structure, however, in contrast to antagonist compound 21, ghrelin or GHRP-6 pushes the side-chain of R283^6.55^ to swing away from the receptor helical core, which induces rotation of TM6 that initiates the cascade of conformational changes in receptor activation (Fig. 4c, d). Concomitantly, the swing of R283^6.55^ leads to the rotameric switches of F279^6.51^ and W276^6.48^, a residue in conserved “micro-switch”. These conformational changes lead to the swing of F272^6.44^ and the pronounced outward displacement of the cytoplasmic end of TM6 (Fig. 4e). The other conserved residues in “micro-switches” (ERY, PIF, and NPxxY) also undergo active-like conformational changes relative to the antagonist-bound receptor and transmit the peptidic agonism signaling to the cytoplasmic end to facilitate receptor-G protein coupling. (Fig. 4f-h). Meanwhile, the conformational changes of F279^6.51^ and W276^6.48^ also cause rotameric switches of F312^7.42^ and Y313^7.43^ and the repacking of the inter-helical hydrophobic contacts between TM6 and TM7, which leads to the inward shift of the cytoplasmic end of TM7 (Fig. 4i).

Our structures also provide clues for understanding the mechanism of the basal activity of ghrelin receptor. The hydrophobic residues F279^6.51^, F312^7.42^, and Y313^7.43^ are closely packed and form a “hydrophobic lock” to packing TM6 and TM7 (Fig. 4i). As aforementioned, these hydrophobic residues primarily transmit peptide agonism signaling with limited impact on peptide binding. This finding raises a hypothesis that this hydrophobic lock may be responsible for the basal activity of the receptor. This hypothesis is further supported by our mutagenesis analysis that substituting F279^6.51^ and F312^7.42^ with alanine dramatically diminishes the basal activity of ghrelin receptor (Extended Data Fig. 9). It should be noted that alanine mutated Y313^7.43^ reduced the surface expression of the receptor. However, when adjusting mutant expression to a comparable WT receptor level, the basal activity displays a similarly significant decrease (Extended Data Fig. 9). Additionally, alanine mutations of E124^3.33^ and R283^6.55^ also lead to the significantly diminished basal activity of the ghrelin receptor, indicating a potential role of the salt bridge formed by these residues in modulating the receptor’s basal activity (Extended Data Fig. 9). Together, the “hydrophobic lock” and the salt bridge may be involved in the regulation of ghrelin receptor’s basal activity.

## Conclusions

Collectively, this study reveals the structural basis for recognition of ghrelin and GHRP-6 by ghrelin receptor and identifies the binding site for the octanoyl group of ghrelin. According to our structure, the octanoyl group is located at cavity II but not cavity I, which is different from the previous modeling studies ^17–19^. With mutagenesis studies, we propose an acyl-modification driven ghrelin-binding model, in which the binding of octanoyl group in cavity II orients the N-terminus of ghrelin to cavity I, and leads to the initiation of signal transduction. Structural comparisons of G_q_-coupled ghrelin receptor bound to ghrelin and GHRP-6 with the antagonist-bound receptor reveal a unique receptor activation mechanism. The binding of peptides causes steric hindrance to push the side-chain of R283^6.55^ swinging away from the helix core and initiates the rotation of TM6. Altogether, these findings enhance our understanding of the molecular basis for acyl-ghrelin recognition and activation of the ghrelin receptor and provide a framework for the drug design targeting the ghrelin receptor.

## Methods

### Construct cloning

The full-length human ghrelin receptor (residues 1-366) was sub-cloned into pFastBac1 vector with an N-terminal haemagglutinin signal peptide (HA) and His × 10 tag followed by BRIL epitope, as well as LgBiT at the C-terminus to facilitate the protein expression and stability. The ghrelin receptor sequence had no additional mutations or loop deletions. The Gα_q_ was designed into a multifunctional chimera based on mini-Gα_s_ skeleton with G_i1_ N-terminus for the binding of Nb35 and scFv16, respectively. Gα_q_, rat Gβ1 with C-terminal HiBiT connected with a 15 residues linker, bovine Gγ2 and Ric8A were cloned into pFastBac1 vector, respectively. All constructs were prepared using homologous recombination (CloneExpress One Step Cloning Kit, Vazyme).

### Protein expression

We used the Bac-to-Bac baculovirus system (ThermoFisher) in *Spodoptera frugiperda* (*sf9*) cells for expression. Cell cultures were grown in ESF 921 serum-free medium (Expression Systems) to a density of 3~4 × 10^6^ cells/ml. Ghrelin receptor, Gα_q_, rat Gβ1, bovine Gγ2 and Ric8A were co-infected at the ratio of 1:1:1:1:1. After infected by 48 hours, the cells were harvested by centrifugation at 2000 rpm (Thermo Fisher, H12000) for 20 minutes and kept frozen at −80 °C for further usage.

### Complex purification

Cell pellets were thawed at R.T. and resuspended in 20 mM HEPES pH 7.5, 100 mM NaCl, 10 mM MgCl_2_, 5 mM CaCl_2_, 2 mM KCl, 0.1 mM TCEP, supplemented with Protease Inhibitor Cocktail (TargetMol, 1 mL/ 100 mL suspension). ScFv16 was applied to stabilize GHRP-6–ghrelin receptor–G_q_ complex, and Nb35 was used to improve the stability of both ghrelin receptor–G_q_ complexes. The monomeric scFv16 and Nb35 were prepared as previously reported ^28,31^. Both ghrelin receptor–G_q_ complexes were formed on the membrane in the presence of 10 μM ligands (ghrelin or GHRP-6, synthesized by GenScript) and treated with apyrase (25 mU/ml, NEB), Nb35-His (15 μg/ml), or scFv16 followed by incubation for 1 hour at R.T. The suspension was then solubilized by 0.5% (w/v) lauryl maltose neopentyl glycol (LMNG, Anatrace) with 0.1% (w/v) cholesteryl hemisuccinate TRIS salt (CHS, Anatrace) for 3 hours. Insoluble material was removed by centrifugation at 65,000g for 40 min and the solubilized complex was incubated overnight at 4 °C with pre-equilibrated Nickel resin (Ni Smart Beads 6FF, SMART Lifesciences) containing 10 mM imidazole. The resin was washed with 15 column volumes of Wash Buffer 1 containing 20 mM HEPES pH 7.5, 100 mM NaCl, 2 mM MgCl_2_, 15 mM imidazole, 0.1% LMNG, 0.02% CHS, 10 μM ligands (ghrelin or GHRP-6) and 15 column volumes of Wash Buffer 2 containing 20 mM HEPES, pH 7.5, 100 mM NaCl, 2 mM MgCl_2_, 25 mM imidazole, 0.01% LMNG, 0.005% GDN (Anatrace), 0.003% CHS, 10 μM ligands (ghrelin or GHRP-6). The complex was then eluted with 5 column volumes of Elution Buffer containing 20 mM HEPES, pH 7.5, 100 mM NaCl, 250 mM imidazole, 2 mM MgCl_2_, 0.01% LMNG, 0.005% GDN, 0.003% CHS and 10 μM ligands (ghrelin or GHRP-6). The complex was concentrated to 0.5 ml using Ultra Centrifugal Filter (ThermoFisher MWCO, 100 kDa) and subjected to size-exclusion chromatography on a Superdex 200 Increase 10/300 column (GE Healthcare) that was pre-equilibrated with 20 mM HEPES pH 7.5, 100 mM NaCl, 2 mM MgCl_2_, 5 μM ligands, 0.00075%(w/v) LMNG, 000025% glyco-diosgenin (GDN, Anatrace) and 0.00015% (w/v) CHS to separate complex from contaminants. Eluted fractions that consisted of receptor and G protein complex were pooled and concentrated to approximately 10 mg/ml for electron microscopy experiments.

### Cryo-EM grid preparation and data collection

For the cryo-EM grids preparation, 3 μl of purified ghrelin-bound complex at 13 mg/ml and GHRP-6-bound ghrelin receptor complex at 8 mg/ml were applied individually onto a glow-discharged holey carbon grid (Quantifoil, Au300 R1.2/1.3) in a Vitrobot chamber (FEI Vitrobot Mark IV). Cryo-EM imaging was performed on a Titan Krios at 300 kV accelerating voltage in the Center of Cryo-Electron Microscopy Research Center, Shanghai Institute of Materia Medica, Chinese Academy of Sciences (Shanghai, China). Micrographs were recorded using a Gatan K3 Summit direct electron detector in counting mode with a nominal magnification of × 81,000, which corresponds to a pixel size of 1.045 Å. Movies were obtained using serialEM at a dose rate of about 26.7 electrons per A^2^ per second with a defocus ranging from −0.5 to −3.0 μm. The total exposure time was 3 s and intermediate frames were recorded in 0.083 s intervals, resulting in an accumulated dose of 80 electrons per A^2^ and a total of 36 frames per micrograph. A total of 5,673 and 3,362 movies were collected for the ghrelin-bound and GHRP-6-bound ghrelin receptor complex, respectively.

### Cryo-EM data processing

Dose-fractionated image stacks for the ghrelin-bound ghrelin receptor–Gα_q_ complex were subjected to beam-induced motion correction using Motion-Cor2.1 ^32^. Contrast transfer function (CTF) parameters for each micrograph were determined by Ctffind4 ^33^. Particle selection, 2D and 3D classifications of the ghrelin-bound ghrelin receptor–G_q_ complex were performed on a binned data set with a pixel size of 2.09Å using RELION-3.0-beta2 ^34^.

For the ghrelin-bound ghrelin receptor–Gα_q_ complex, semi-automated particle selection yielded 4,598,528 particle projections. The projections were subjected to 2D classification to discard particles in poorly defined classes, producing 1,842,606 particle projections for further processing. The map of the D1R–Gs complex low-pass filtered to 40 Å was used as a reference model for four rounds of maximum-likelihood-based 3D classifications, resulting in one well-defined subset with 912,636 projections. A map generated by 3D refinement was subsequently post-processed in DeepEMhancer ^35^. The final refinement generated a map with an indicated global resolution of 2.9 Å at a Fourier shell correlation of 0.143.

For the GHRP-6-bound ghrelin receptor–Gα_q_ complex, semi-automated particle selection yielded 2,728,266 particle projections. The projections were subjected to 2D classification, producing 1,523,752 particle projections for further processing. The map of the D1R–G_s_ complex low-pass filtered to 40 Å was used as a reference model for three rounds of maximum-likelihood-based 3D classifications, resulting in two well-defined subsets with 262,892 projections. A map generated by 3D refinement was subsequently post-processed in DeepEMhancer ^35^. The final refinement generated a map with an indicated global resolution of 3.2 Å at a Fourier shell correlation of 0.143. Local resolution for both density maps was determined using the Bsoft package with half maps as input maps ^36^.

### Model building and refinement

The crystal structure of the ghrelin receptor (PDB: 6KO5) was used as an initial model for model rebuilding and refinement against the electron microscopy maps of ghrelin receptor–Gα_q_ complexes. The structure of the G_q_ part of the 5-HT_2A_ complex (PDB: 6WHA) was used as initial models for model building of the ghrelin/GHRP-6 bound ghrelin receptor–Gα_q_–Nb35–(scFv16) complex. The initial models were docked into the electron microscopy density maps using Chimera ^37^ followed by iterative manual adjustment and rebuilding in COOT ^38^. Real-space refinement and reciprocal space refinement were performed using Phenix programs ^39^. The model statistics were validated using MolProbity ^40^. Structure figures were prepared in Chimera and PyMOL (https://pymol.org/2/). The final refinement statistics are provided in Extended Data Table 1. The extent of any model overfitting during refinement was measured by refining the final model against one of the half-maps and by comparing the resulting map versus model FSC curves with the two half-maps and the full model.

### Ligand-binding assays

Ligand binding was performed with a homogeneous time-resolved fluorescence-based assay. N-terminal-SNAP-tagged ghrelin receptor (WT or with various mutations) and full-length ghrelin labeled with the dye A2 on an additional cysteine at the C-terminal end of the peptide (ghrelin-A2, synthesized by Vazyme, China) were used as previously described ^41^.

HEK293 cells transfected with SNAP-ghrelin receptor (WT or mutants) were seeded at a density of 1 × 10^6^ cells into 3 cm dish and incubated for 24 hours at 37 °C in 5% CO_2_. Cell culture medium was removed and Tag-lite labeling medium with 100 nM of SNAP-Lumi4-Tb (Cisbio, SSNPTBC) was added, and the cells were further incubated for 1 hour at 37 °C in 5% CO_2_. The excess of SNAP-Lumi4-Tb was then removed by washing 4 times with 1 ml of Tag-lite labeling medium.

For saturation binding experiments, we incubated cells with increasing concentrations of ghrelin-A2 in the presence or absence of 10 μM unlabeled ghrelin for 1 hour at R.T. Signal was detected using the Multimode Plate Reader (PerkinElmer EnVision) equipped with an HTRF optic module allowing a donor excitation at 340 nm and a signal collection both at 665 nm and at 620 nm. HTRF ratios were obtained by dividing the acceptor signal (665 nm) by the donor signal (620 nm). Kd values were obtained from binding curves using Prism 5.0 software (GraphPad Software).

### Calcium assay

The wild-type ghrelin receptor gene was subcloned in the pcDNA3.0 vector with an N-terminal HA signal peptide. Mutations were introduced by QuickChange PCR. All of the constructs were verified by DNA sequencing. HEK293 cells transfected with HA-tagged WT ghrelin receptor or mutants were seeded at a density of 4× 10^4^ cells per well into 96-well culture plates and incubated for 24 hours at 37 °C in 5% CO2.The cells were then incubated with 2 μmol/L Fluo-4 AM in HBSS (5.4 mmol/L KCl, 0.3 mmol/L Na2HPO4, 0.4 mmol/L KH2PO4, 4.2 mmol/L NaHCO3, 1.3 mmol/L CaCl_2_, 0.5 mmol/L MgCl_2_, 0.6 mmol/L MgSO4, 137 mmol/L NaCl, 5.6 mmol/L D-glucose and 250 μmol/L sulfinpyrazone, pH 7.4) at 37 °C for 40 min. After thorough washing, 50 μL of HBSS was added. After incubation at R.T. for 10 min, 25 μL of agonist was dispensed into the well using a FlexStation III microplate reader (Molecular Devices), and the intracellular calcium change was recorded at an excitation wavelength of 485 nm and an emission wavelength of 525 nm. EC_50_ and *E_max_* values for each curve were calculated by Prism 5.0 software.

### Inositol phosphate accumulation assay

IP1 production was measured using the IP-One HTRF kit (Cisbio, 621PAPEJ) as described previously ^41^. Briefly, 24 hours after transfection, cells were harvested and re-suspended in PBS at a density of 4 × 10^6^ cells/ml. Cells were then plated onto 384-well assay plates at 20,000 cells/5 μl/well. Another 5 μl IP1 stimulation buffer containing ligand was added to the cells, and the incubation lasted for 30 min at R.T. As a negative control, cells transfected with pcDNA3.0 empty vector were also tested. Intracellular IP1 measurement was carried with the IP-One HTRF kit and EnVision multiplate reader according to the manufacturer’s instructions. The HTRF ratio was converted to a response (%) using the following formula: response (%) = (ratio of sample – ratio of the negative control)/ (ratio of WT – ratio of the negative control) ×100.

### Cell-surface expression assay

Cell-surface expression for each mutant was monitored by a fluorescence-activated cell sorting (FACS) assay. In brief, the expressed cells were incubated with mouse anti-HA-FITC antibody (Sigma) for 20 min at 4 °C, and then a 9-fold excess of PBS was added to cells. Finally, the surface expression of the ghrelin receptor was monitored by detecting the fluorescent intensity of FITC using a BD ACCURI C6.

### Statistical analysis

All functional study data were analyzed using Prism 8 (GraphPad) and presented as means ± S.E.M. from at least three independent experiments. Concentration-response curves were evaluated with a three-parameter logistic equation. EC_50_ is calculated with the Sigmoid three-parameter equation. The significance was determined with two-side, one-way ANOVA with Tukey’s test, and *P* < 0.05 *vs*. wild-type (WT) was considered statistically significant.

## Acknowledgements

The Cryo-EM data were collected at Cryo-Electron Microscopy Research Center, Shanghai Institute of Material Medica. We thank the staff of the National Center for Protein Science (Shanghai) Electron Microscopy facility for instrument support. This work was partially supported by the National Natural Science Foundation (31770796 to Y.J., 81730099 to X.X.), the National Science and Technology Major Project (2018ZX09711002-002-002) to Y.J., the National Key R&D Programs of China (2018YFA0507002), the Shanghai Municipal Science and Technology Major Project (2019SHZDZX02), the CAS Strategic Priority Research Program (XDB37030103) to H.E.X., and the Shanghai Municipal Science and Technology Commission Grant (20S11903200) to X. X.

## Author Contributions

Y. J., H.E.X. and X.X conceived, designed and supervised the overall project. Y.J., H.E.X., X.X., Y.W., S.-M.G. Y.-W.Z., and P.-Y.X. participated in data analysis and interpretation; Y.W. generated the ghrelin receptor insect cell expression constructs, established the ghrelin–ghrelin receptor–G_q_ – Nb35 and GHRP-6–ghrelin receptor–G_q_–Nb35–scFv16 complex purification protocol and prepared samples for the cryo-EM as well as data collection; S.-M.G. and Y. Y. designed all of the mutants for the ligand-binding pockets and executed all the cellular experiments including the ligand-binding assay, Calcium assay and IP1 assay supervised by X.X.; Y.-W.Z. performed cryo-EM data processing and P.-Y.X. performed model building and refinement; X.-H.H. modelled disease-related mutants; J.G. assisted Y.W. with protein complex purification and sample preparation for cryo-EM; W.C.Y. designed G_q_ protein constructs; Y.J., Y.W., S.-M.G. prepared the figures; Y.J., H.E.X., X.X., Y.W., S.-M.G. wrote the manuscript with input from all authors.

## Data availability

All data is available in the main text or the supplementary materials. Materials are available from the corresponding authors upon reasonable request.

---

**Extended Data Fig. 1.**
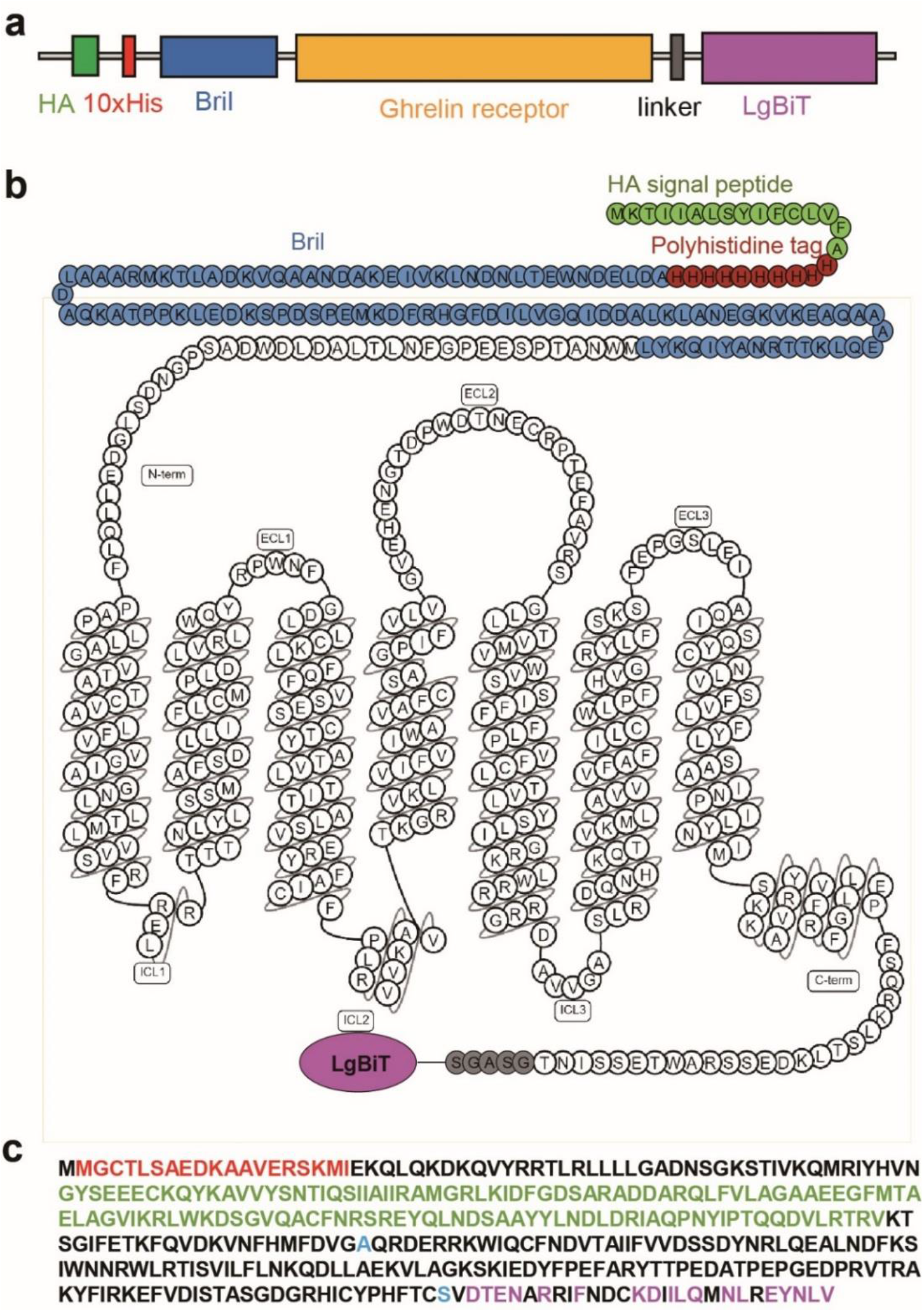
Ghrelin receptor construct and Gα_q_ chimera/mutant constructs used in this study. **a, b**, Schematic representation (**a**) and snake model (**b**) of the ghrelin receptor construct. **c**, Sequence of engineered Gα_q_, the skeleton is based on miniGα_s_ (for Nb35 binding), which is shown in black. N-terminus in red is replaced by Gα_i1_ (for scFv16 binding). Two dominant-negative mutations are shown in cyan, WT Gα_q_ is colored in magenta.

**Extended Data Fig. 2.**
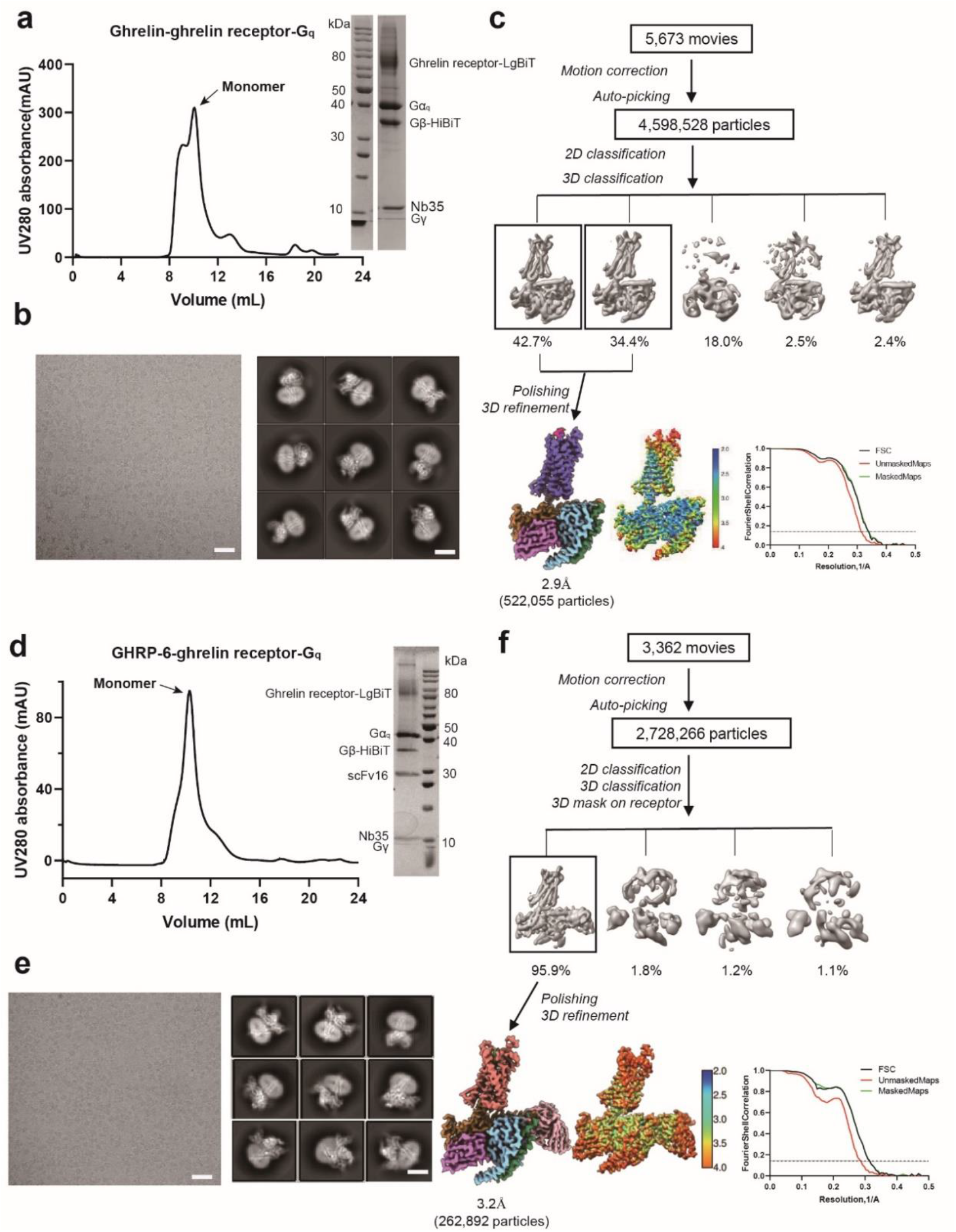
Ghrelin receptor–G_q_ complex purification and cryo-EM data processing. **a**, Representative elution profile of His-purified ghrelin–ghrelin receptor–LgBiT–G_q_ complex and SDS–PAGE of the size-exclusion chromatography peak. **b**, Cryo-EM micrographs of ghrelin–ghrelin receptor–LgBiT–G_q_ complex (scale bar: 50 nm) and 2D class averages (scale bar: 5 nm). **c**, Flow chart of the cryo-EM data processing for the ghrelin–ghrelin receptor–LgBiT–G_q_ complex. **d**, Representative elution profile of His-purified GHRP-6–ghrelin receptor–LgBiT–G_q_ complex and SDS–PAGE of the size-exclusion chromatography peak. **e**, Cryo-EM micrographs of GHRP-6–ghrelin receptor–LgBiT–G_q_ complex (scale bar: 50 nm) and 2D class averages (scale bar: 5 nm). **f**, Flow chart of the cryo-EM data processing for the GHRP-6–ghrelin receptor–LgBiT–G_q_ complex.

**Extended Data Fig. 3.**
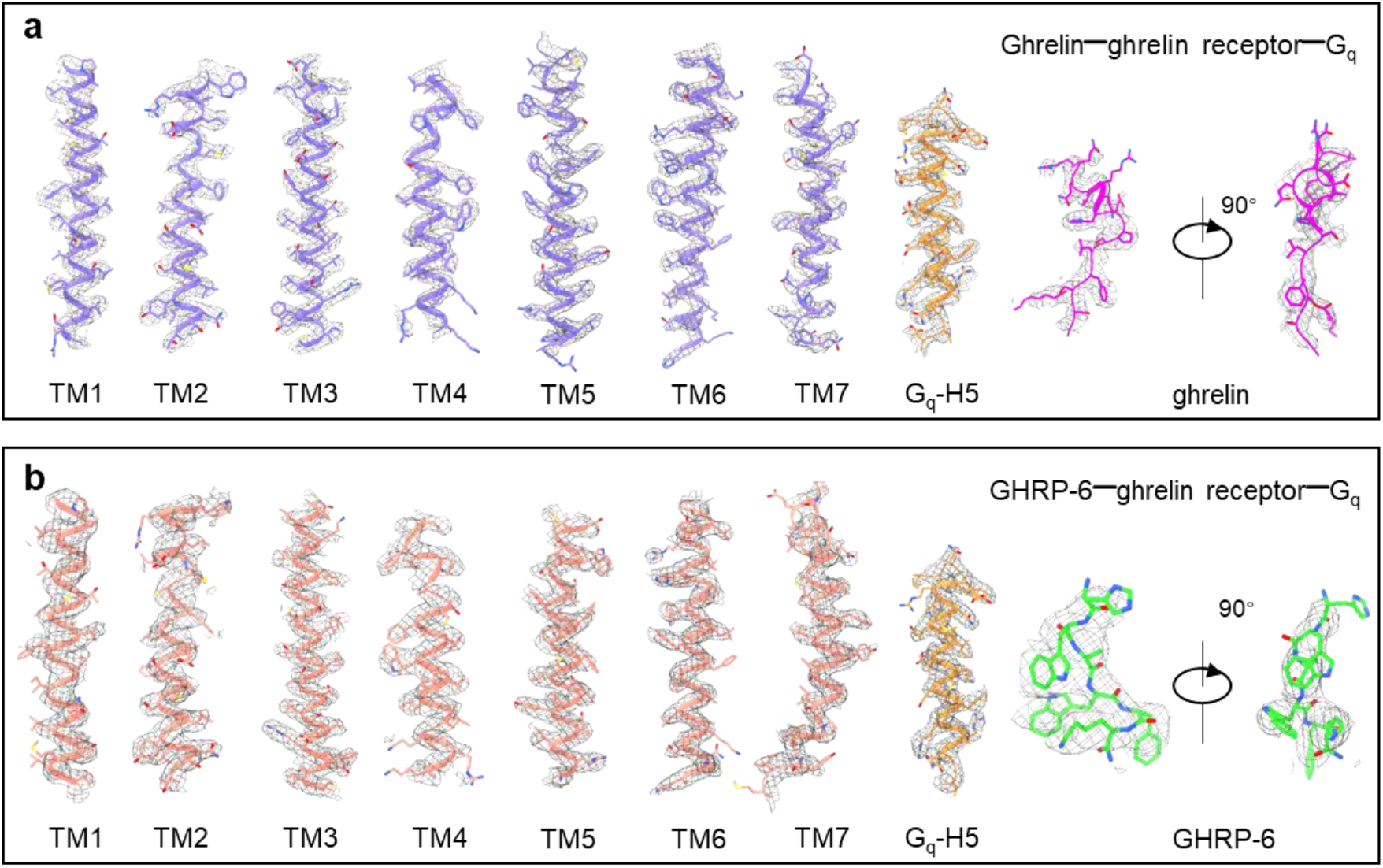
Overall resolution analysis of electron density of transmembrane helices, α5-helix of G_q_, ghrelin and GHRP-6. **a**, EM density and model of all transmembrane helices of ghrelin-bound ghrelin receptor. **b**, GHRP-6**-**bound ghrelin receptor and α5 helix of Gα_q_ subunit.

**Extended Data Fig. 4.**
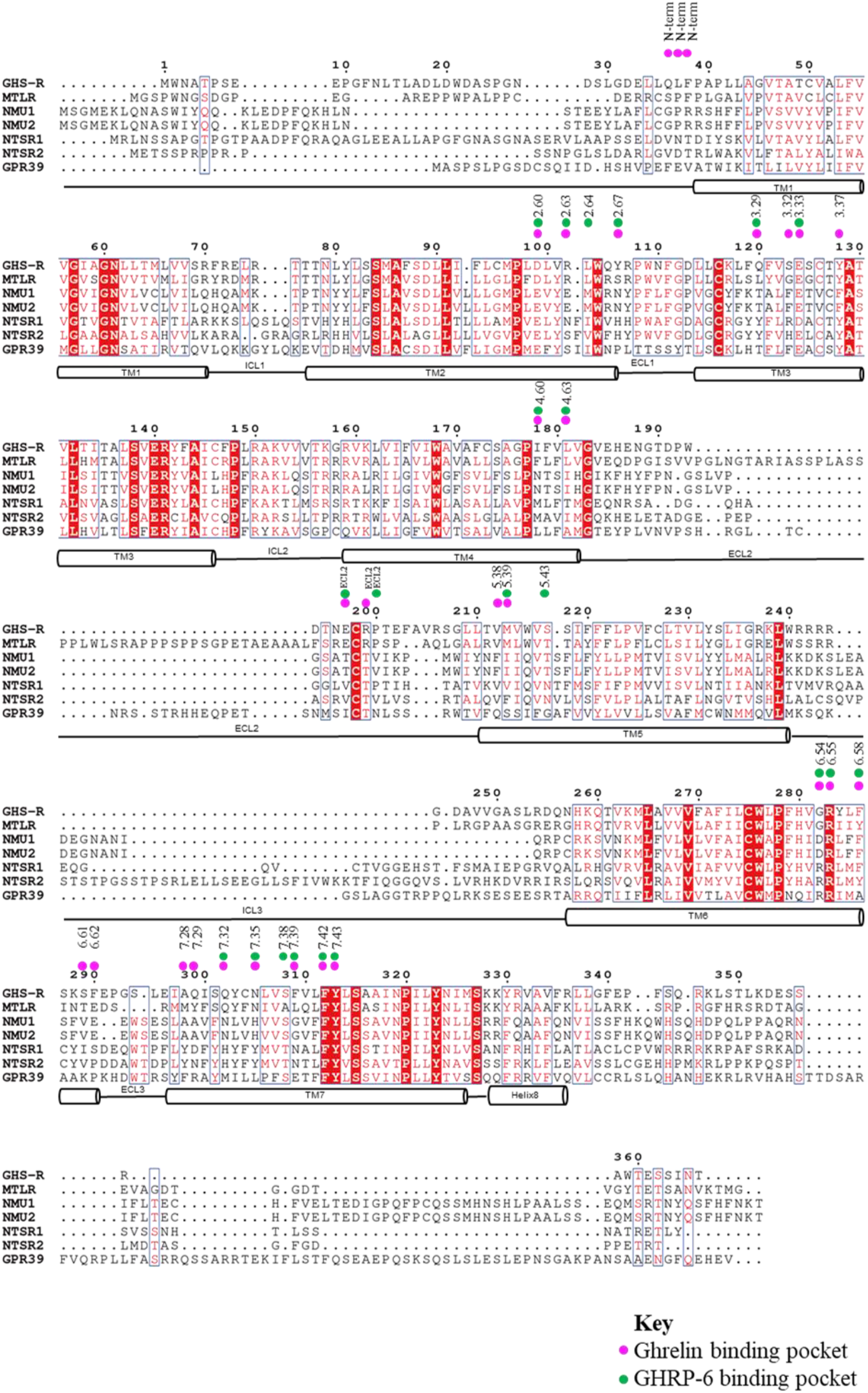
Sequence alignment of the ghrelin receptor family. The sequences shown are those for motilin receptor (MTLR), neuromedin U receptor 1/2 (NMU1/2), neurotensin receptors 1/2 (NTSR1/2) and orphan receptor (GPR39), The sequence alignment was created using Clustalw and ESPript 3.0 servers. α-helices for ghrelin receptor are shown as columns. The binding-pocket residues are shown as magenta (ghrelin) and green (GHRP-6) dots.

**Extended Data Fig. 5.**
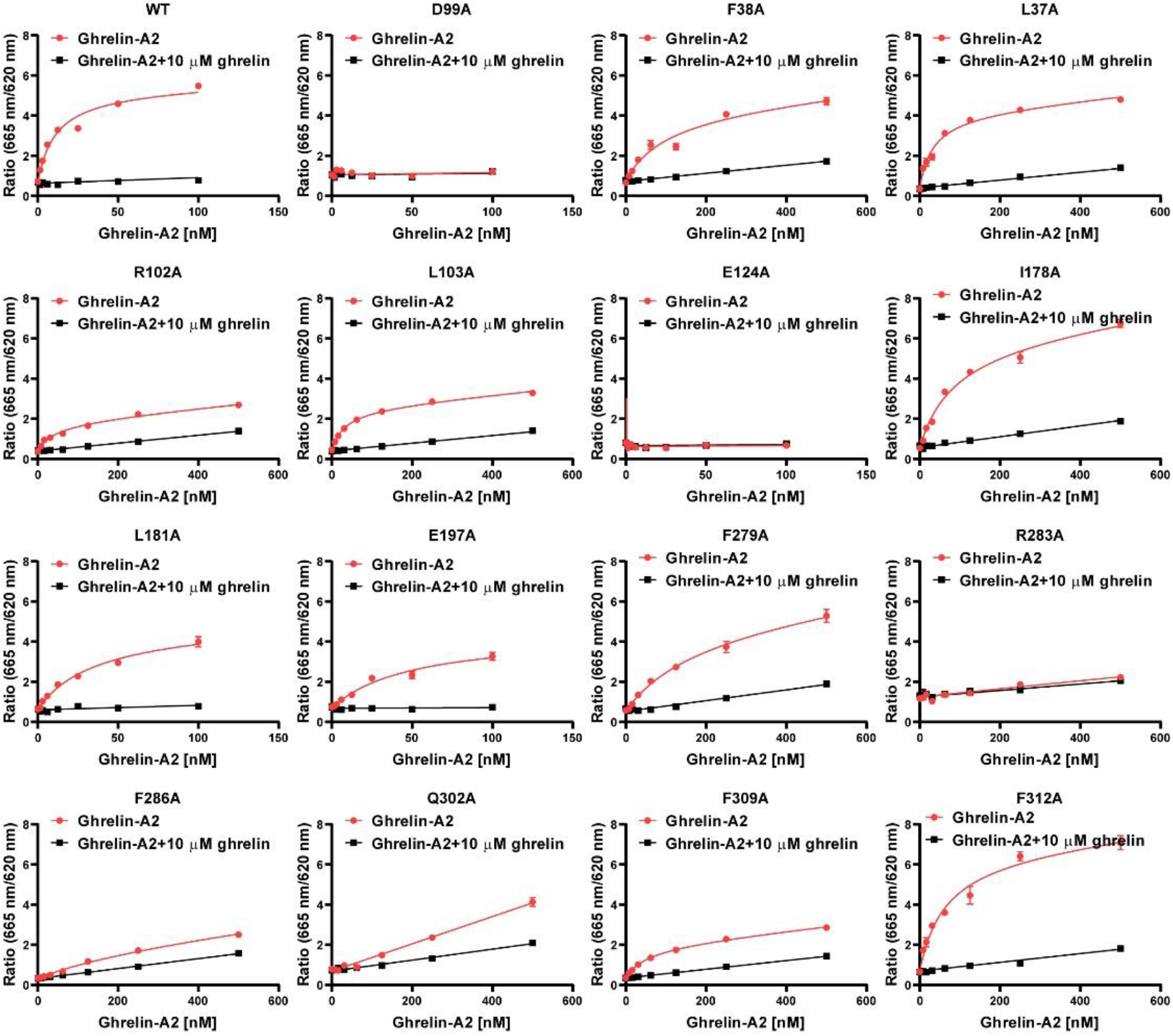
Saturation experiments of ghrelin-A2 binding to the WT and mutant ghrelin receptor. Effects of different mutations within the ligand-binding pocket of ghrelin receptor. Amino acids located in the ghrelin binding pocket were mutated to an alanine residue, saturation binding experiments on HEK293 cells transfected with different ghrelin receptor mutations. Each point represents mean ± S.E.M. from three or four independent experiments.

**Extended Data Fig. 6.**
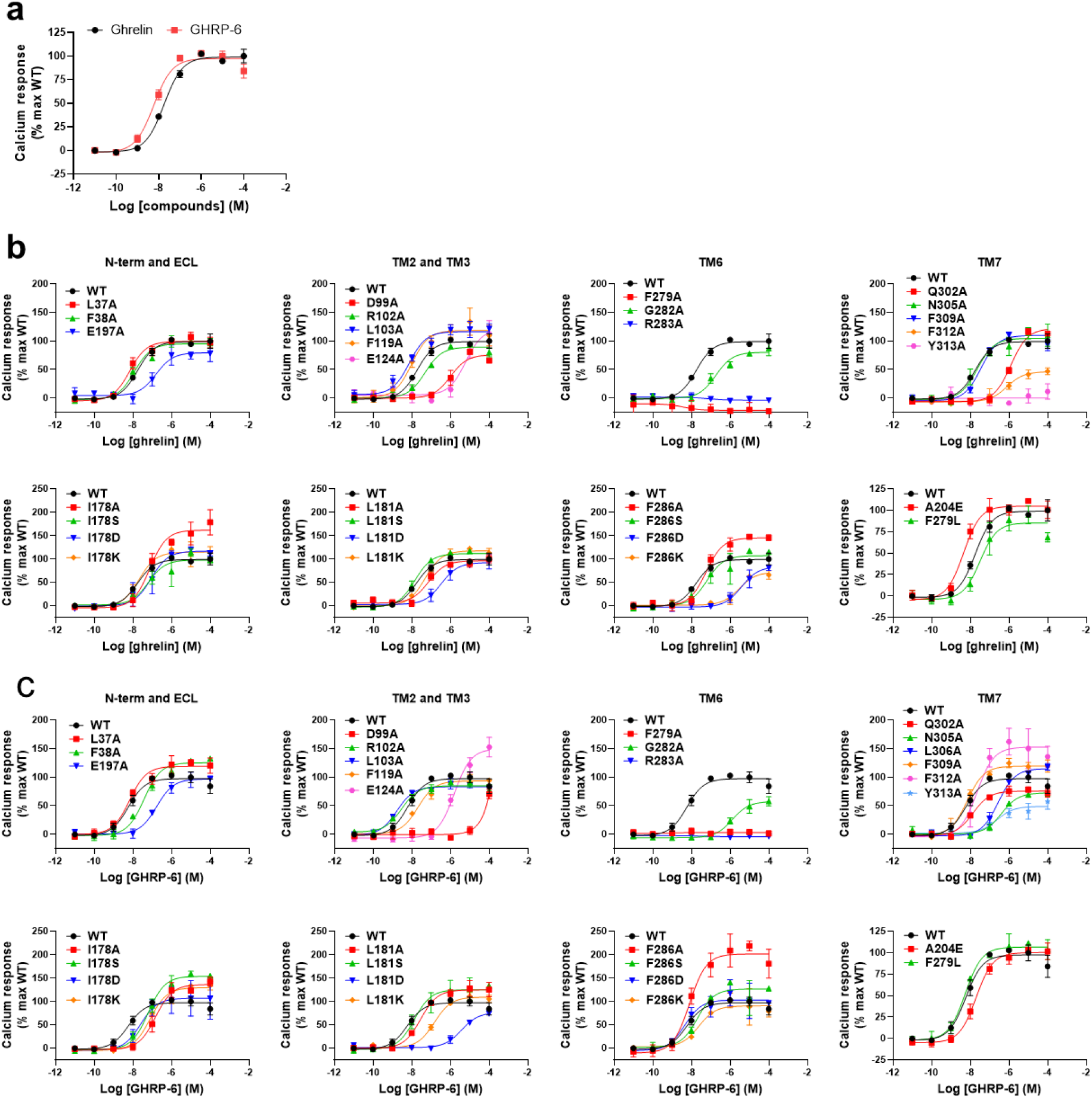
Ghrelin/GHRP-6 response curves for WT or ghrelin receptor mutants. HEK293 cells were transfected with WT or ghrelin receptor mutant constructs. Intracellular calcium signals were monitored after stimulation with ghrelin or GHRP-6. Each point represents mean ± S.E.M. from three independent experiments. **a**, Comparison of the activities of ghrelin or GHRP-6 on WT ghrelin receptor. GHRP-6 displays comparable potency with ghrelin for ghrelin receptor activation. **b**, Effects of ghrelin receptor mutations on ghrelin induced calcium mobilization. **c**, Effects of ghrelin receptor mutations on GHRP-6 induced calcium mobilization.

**Extended Data Fig. 7.**
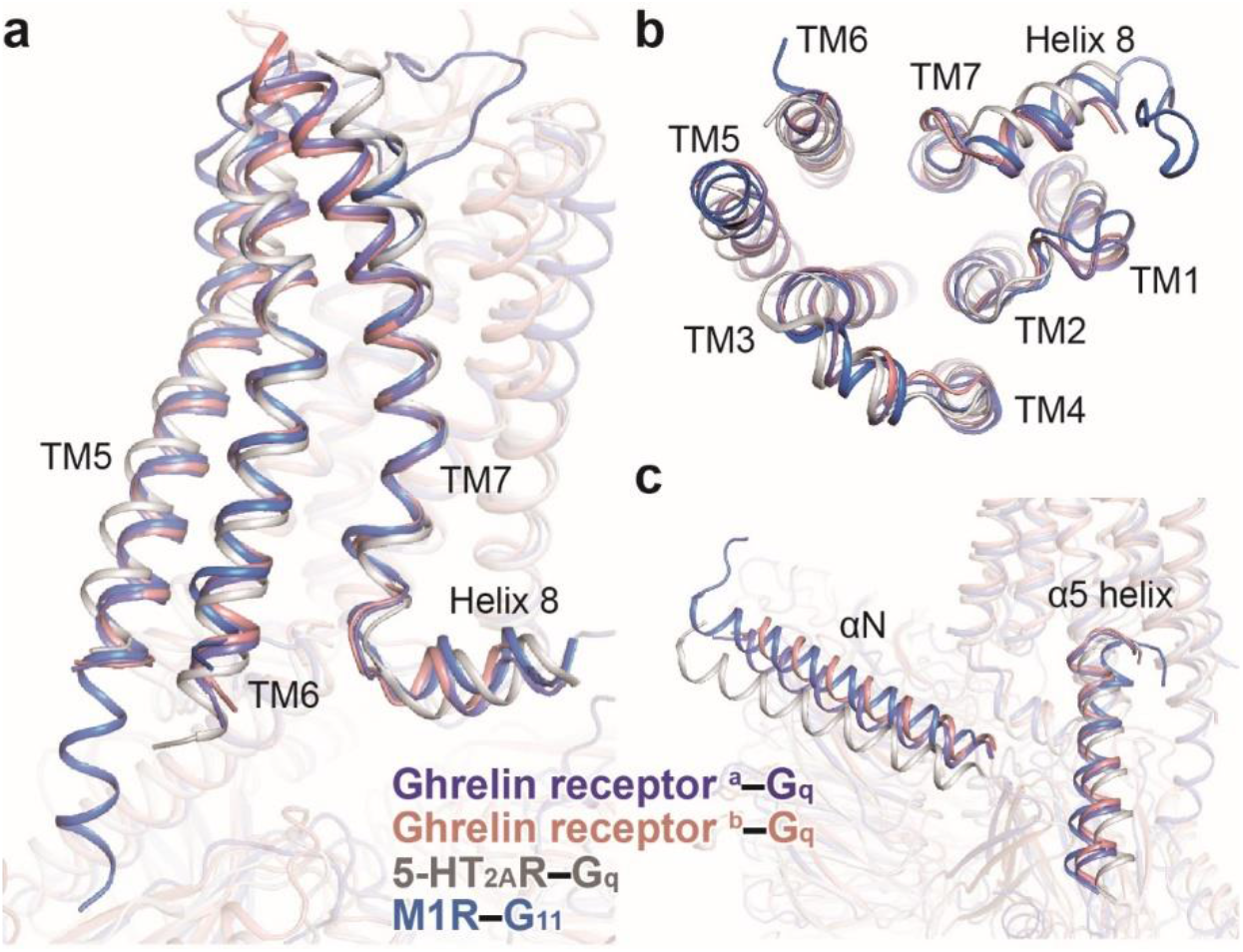
Structural comparison of G_q_-coupled ghrelin receptors with other G_q/11_-coupled GPCRs solved to date. **a, b**, Structural superpositions of G_q/11_-coupled receptors. Orthogonal view (**a**), extracellular view (**b**). Structures of TMs 5-7 from receptors are highlighted. **c**, Conformational comparison of αN and α5 helix of G_q/11_ proteins. αN, N-terminus of Gα_q_ subunit. ^a^, ghrelin-bound ghrelin receptor complex (slate blue). ^b^, GHRP-6-bound ghrelin receptor complex (salmon). G_q_-coupled 5-HT_2A_R is shown in grey (PDB: 6WHA), and G_11_-coupled M1R is displayed in marine (PDB: 6OIJ). The Ligands are omitted for clear presentation.

**Extended Data Fig. 8.**
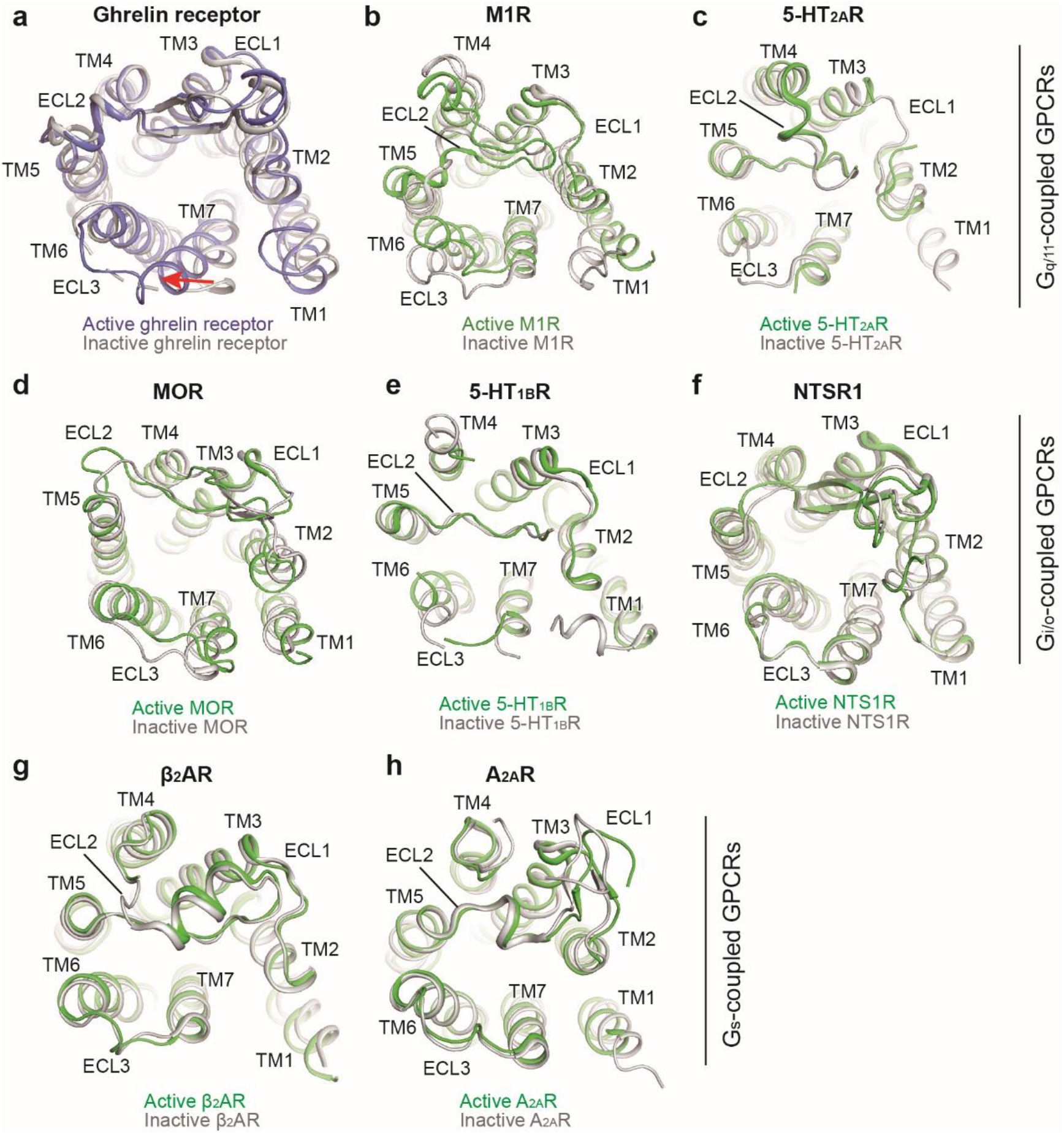
Conformational changes of the extracellular end of TM7 across representative class A GPCRs. Structural superposition of G_q_/11-coupled (**a-c**), G_i/o_-coupled (**d-f**), and Gs-coupled GPCRs (**g, h**) in the extracellular view. **a**, active ghrelin receptor (ghrelin-bound) and antagonist-bound ghrelin receptor; **b**, Active M1R (PDB: 6OIJ) and inactive M1R (PDB: 6WJC); **c**, Active 5-HT_2A_R (PDB: 6WHA) and inactive 5-HT_2A_R (PDB: 6A94); **d**, Active MOR (PDB: 6DDE) and inactive MOR (PDB: 4DKL); **e**, Active 5-HT_1B_R (PDB: 5G79) and inactive 5-HT_1B_R (PDB: 5V54); **f**, Active NTSR1 (PDB: 4GRV) and inactive NTSR1 (PDB: 4BUO); **g**, Active β_2_AR (PDB: 3SN6) and inactive β_2_AR (PDB: 3NYA); **h**, Active A_2A_R (PDB: 5G53) and inactive A_2A_R (PDB: 3EML). Except for active ghrelin receptor (slate blue), other active receptors are colored in green, and all inactive or antagonist-bound receptors are shown in grey.

**Extended Data Fig. 9.**
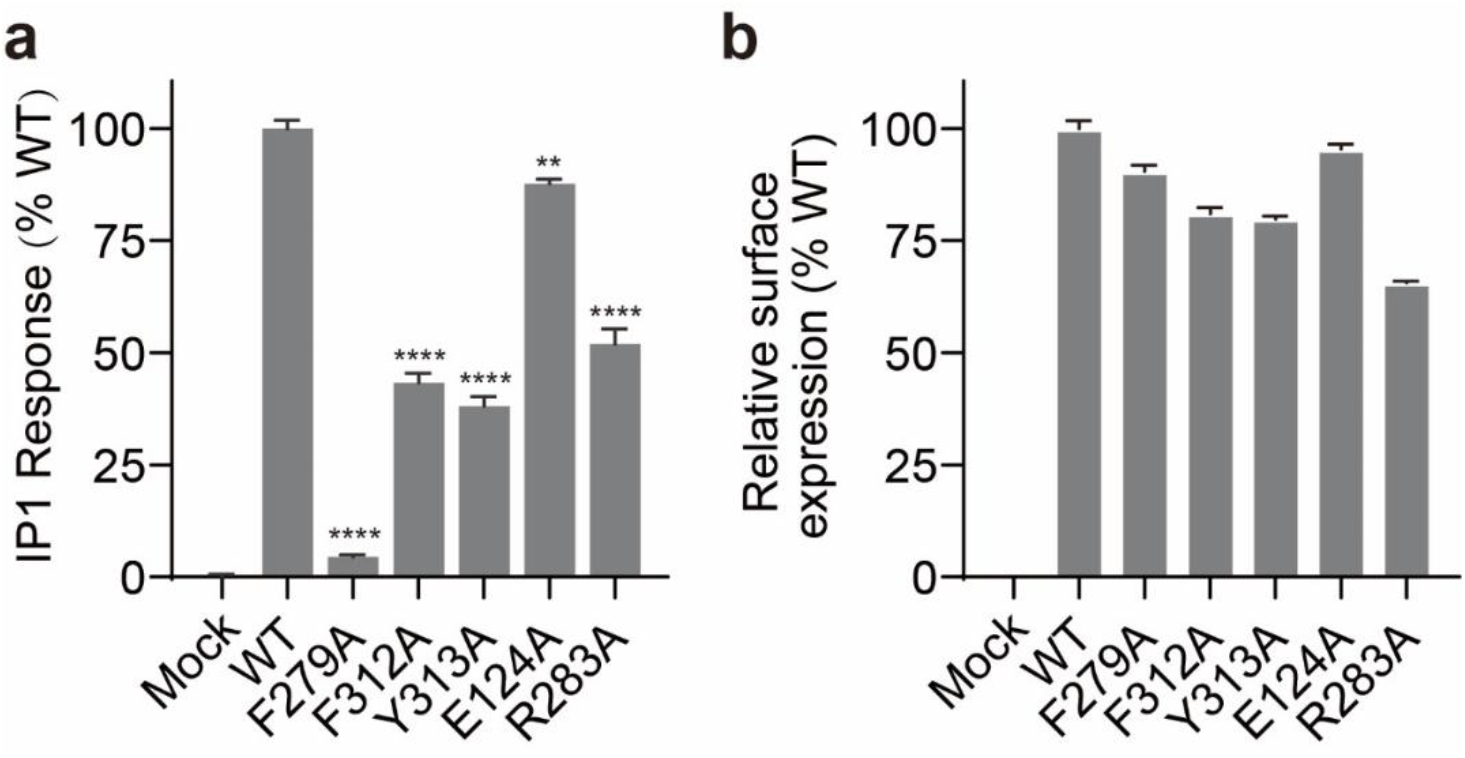
Effects of alanine mutations of residues on the basal activity of the ghrelin receptor. Inositol phosphate accumulation assay was performed to evaluate the basal activity of the ghrelin receptor (**a**). Residues in the “hydrophobic lock” (F279^6.51^, F312^7.42^, and Y313^7.43^) and residues forming a salt bridge (E124^3.33^ and R283^6.55^) were substituted with alanines. The cell surface expression of these mutants was determined by using flow cytometry (**b**). Considering the Y313A mutation decreases the cell surface expression, a 4-fold amount of the Y313A construct relative to other mutants was transfected into HEK-293 cells to achieve a comparable expression level compared with wild-type (WT) receptor.

**Extended Data Table 1.**
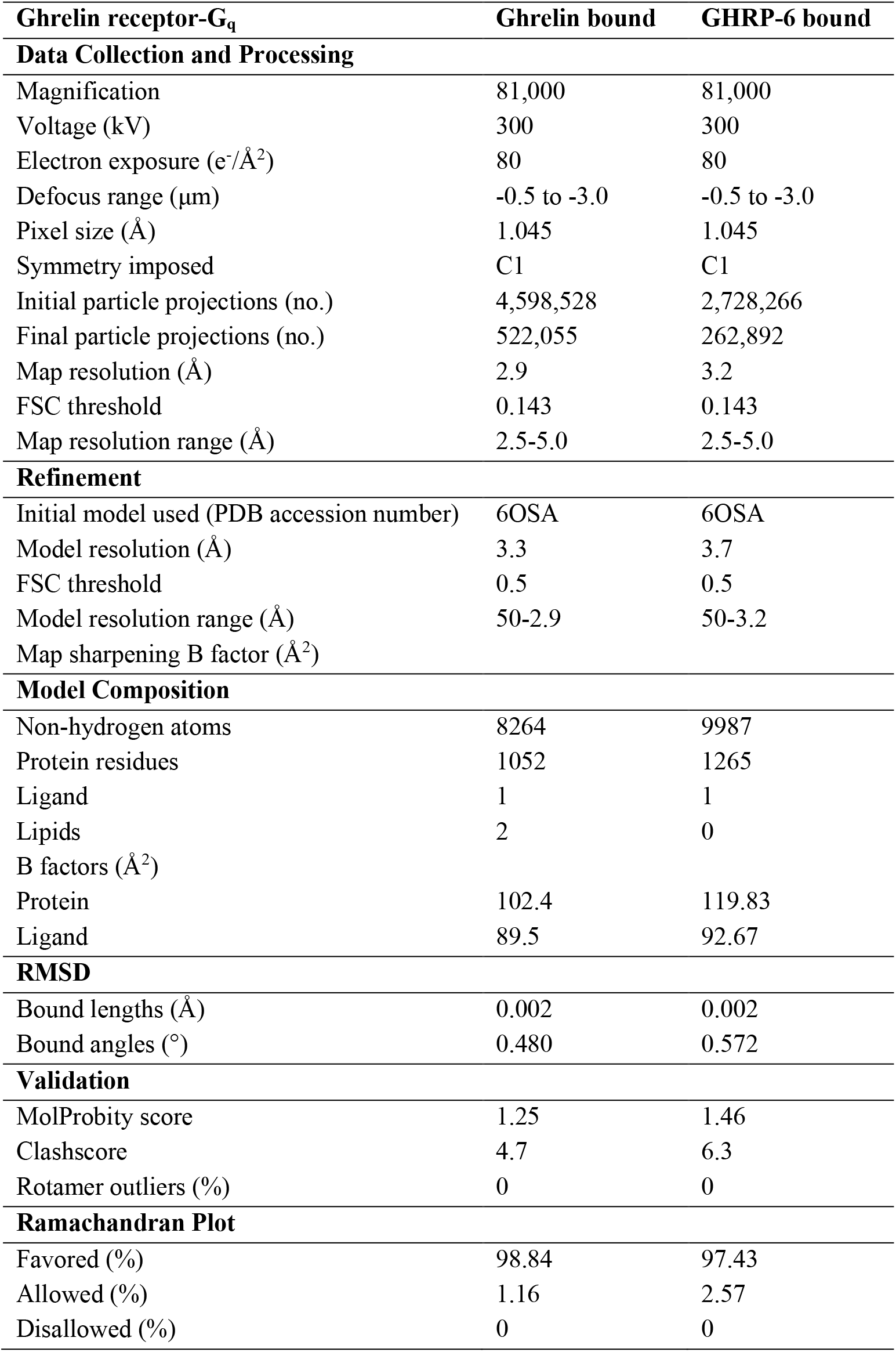
Cryo-EM data collection, model refinement, and validation statistics.

| Ghrelin receptor-G<sub>q</sub> | Ghrelin bound | GHRP-6 bound |
| --- | --- | --- |
| **Data Collection and Processing** |  |  |
| Magnification | 81,000 | 81,000 |
| Voltage (kV) | 300 | 300 |
| Electron exposure (e<sup>-</sup>/Å<sup>2</sup>) | 80 | 80 |
| Defocus range (μm) | -0.5 to -3.0 | -0.5 to -3.0 |
| Pixel size (Å) | 1.045 | 1.045 |
| Symmetry imposed | C1 | C1 |
| Initial particle projections (no.) | 4,598,528 | 2,728,266 |
| Final particle projections (no.) | 522,055 | 262,892 |
| Map resolution (Å) | 2.9 | 3.2 |
| FSC threshold | 0.143 | 0.143 |
| Map resolution range (Å) | 2.5-5.0 | 2.5-5.0 |
| **Refinement** |  |  |
| Initial model used (PDB accession number) | 6OSA | 6OSA |
| Model resolution (Å) | 3.3 | 3.7 |
| FSC threshold | 0.5 | 0.5 |
| Model resolution range (Å) | 50-2.9 | 50-3.2 |
| Map sharpening B factor (Å<sup>2</sup>) |  |  |
| **Model Composition** |  |  |
| Non-hydrogen atoms | 8264 | 9987 |
| Protein residues | 1052 | 1265 |
| Ligand | 1 | 1 |
| Lipids | 2 | 0 |
| B factors (Å<sup>2</sup>) |  |  |
| Protein | 102.4 | 119.83 |
| Ligand | 89.5 | 92.67 |
| **RMSD** |  |  |
| Bound lengths (Å) | 0.002 | 0.002 |
| Bound angles (°) | 0.480 | 0.572 |
| **Validation** |  |  |
| MolProbity score | 1.25 | 1.46 |
| Clashscore | 4.7 | 6.3 |
| Rotamer outliers (%) | 0 | 0 |
| **Ramachandran Plot** |  |  |
| Favored (%) | 98.84 | 97.43 |
| Allowed (%) | 1.16 | 2.57 |
| Disallowed (%) | 0 | 0 |

**Extended Data Table 2.** *Kd* and *B_max_* of ghrelin-A2 binding to ghrelin receptor mutants and EC_50_ values of different ligands on ghrelin receptor mutants.

|  | Ligand-binding assays |  |  | Calcium assay |  |  |  | Cell-surface expression |
| --- | --- | --- | --- | --- | --- | --- | --- | --- |
|  | ghrelin-A2 |  |  | ghrelin |  | GHRP-6 |  |  |
| | $K_d$ (nM) | fold of change | $B_{max}$ (% max WT) | $EC_{50}$ (nM) | fold of change | $EC_{50}$ (nM) | fold of change | Relative to WT |
| WT | 10.84±0.19 | 1 | 100 | 18.0±1.2 | 1 | 5.97±0.6 | 1 | 100±1.9 |
| L37<sup>N-term</sup>A | 32.9±0.44 | 3 | 80.8 | 7.33±1.43 | 0.4 | 6.94±1.16 | 1.2 | 98.43±0.8 |
| F38<sup>N-term</sup>A | 99.27±16.5 | 9.2 | 77.1 | 12.6±2.32 | 0.7 | 39.9±6.92 | 6.7 | 93.64±0.5 |
| D99<sup>2.60</sup>A | UD | +++ | UD | 1121.3±335.1 | 62.3 | UD | +++ | 95.09±1.9 |
| R102<sup>2.63</sup>A | 41.9±2.33 | 3.9 | 30.8 | 47.6±6.78 | 2.6 | 2.86±0.33 | 0.5 | 96.7±2.5 |
| L103<sup>2.64</sup>A | 30.9±0.23 | 2.9 | 45.7 | 7.02±2.28 | 0.4 | 1.62±0.11 | 0.3 | 99.11±2.3 |
| E124<sup>3.33</sup>A | UD | +++ | UD | 6547±2390 | 363.7 | 1646±426 | 275.7 | 93.73±1.3 |
| I178<sup>4.60</sup>A | 84.30±1.87 | 7.8 | 117.5 | 76.1±6.9 | 4.2 | 154.7±23.5 | 25.8 | 94.6±1.9 |
| I178<sup>4.60</sup>S | NT | NT | NT | 67.2±11.1 | 3.7 | 63±7.2 | 10.6 | 100.18±1 |
| I178<sup>4.60</sup>D | NT | NT | NT | 96.4±21.3 | 5.4 | 25.4±1.8 | 4.3 | 88.05±0.8 |
| I178<sup>4.60</sup>K | NT | NT | NT | 22.5±4 | 1.3 | 79.2±5.7 | 13.3 | 93.22±2.1 |
| L181<sup>4.63</sup>A | 37.75±2.5 | 3.5 | 91 | 75.2±7 | 4.2 | 29.9±3.8 | 5 | 99.53±3.5 |
| L181<sup>4.63</sup>S | NT | NT | NT | 14.9±1.5 | 0.8 | 16.9±5.8 | 2.8 | 95.13±1.2 |
| L181<sup>4.63</sup>D | NT | NT | NT | 368.5±59.9 | 20.4 | 3854±512.6 | 64.5 | 97.24±1 |
| L181<sup>4.63</sup>K | NT | NT | NT | 50.1±3.5 | 2.8 | 151.1±11.8 | 25.3 | 89.91±1.1 |
| E197<sup>ECL2</sup>A | 50.88±0.25 | 4.7 | 81.4 | 161.7±33.5 | 8.9 | 173±9.9 | 29 | 103.8±1.7 |
| F279<sup>6.51</sup>A | 180.1±23.7 | 16.7 | 98.1 | UD | +++ | UD | +++ | 101.27±1.9 |
| G282<sup>6.54</sup>A | NT | NT | NT | 225±93.0 | 12.5 | 1387.6±352.4 | 232.3 | 90.13±3.5 |
| R283<sup>6.55</sup>A | UD | +++ | UD | UD | +++ | UD | +++ | 63.24±1.8 |
| F286<sup>6.58</sup>A | 275±27.5 | 25.5 | 32.7 | 57.9±2.8 | 3.2 | 8.1±0.9 | 1.4 | 91.26±0.9 |
| F286<sup>6.58</sup>S | NT | NT | NT | 75.2±21.1 | 4.2 | 21.2±2.7 | 3.6 | 99.6±0.6 |
| F286<sup>6.58</sup>D | NT | NT | NT | 4661±1415 | 258.9 | 4.6±0.5 | 0.8 | 95.36±1 |
| F286<sup>6.58</sup>K | NT | NT | NT | 2966±46.1 | 164.8 | 24.6±3.3 | 4.1 | 101.09±1 |
| S289<sup>6.61</sup>A | NT | NT | NT | 19.9±6.41 | 1.1 | 1.89±0.44 | 0.3 | 99.84±1.5 |
| Q302<sup>7.32</sup>A | >500 | +++ | UD | 1267±48.7 | 70.3 | 13.7±0.5 | 2.3 | 92.67±3 |
| N305<sup>7.35</sup>A | NT | NT | NT | 24.6±7.43 | 1.4 | 420±50.5 | 70.4 | 85.3±1.2 |
| F309<sup>7.39</sup>A | 52.5±0.85 | 4.9 | 34.4 | 36.5±4.25 | 2 | 7.01±0.82 | 1.2 | 79.97±1.5 |
| F312<sup>7.42</sup>A | 64.47±8.60 | 6 | 128.8 | 794±148 | 43.8 | 23±2.9 | 3.9 | 82±2.3 |
| Y313<sup>7.43</sup>A | NT | NT | NT | UD | +++ | 326±160 | 54.6 | 25.14±0.8 |
| A204<sup>ECL2</sup>E | NT | NT | NT | 4.67±0.59 | 0.3 | 21.1±3.60 | 3.5 | 78.78±1.1 |
| F279<sup>6.51</sup>L | NT | NT | NT | 41.4±16.3 | 2.3 | 5.51±0.39 | 0.9 | 99.52±1 |

UD, Undetectable; +++, Exceedingly high fold of change due to the undetectable or very low agonist activity; NT, Not tested.

